# Tunable Tau Expression in *C. elegans* Neurons Reveals that Early-AD Tau Phosphorylation Selectively Impacts Behavior and Mitochondrial Quality Control

**DOI:** 10.64898/2026.01.26.701793

**Authors:** T Carroll, D Pfendler, H Alhaj Arhayem, R Thoma, A Müller-Eigner, A. Straut, G.V.W Johnson, K Nehrke

## Abstract

Tau protein accumulates myriad post-translational modifications as Alzheimer’s disease (AD) progresses, and early-disease tau modifications such as phosphorylation at threonine 231 (T231) likely play a key role in AD pathogenesis. Here, a series of “tunable tau” strains was developed in *C. elegans* to test the relative impact of tau pseudo-phosphorylation of T231 (T231E) compared to protein expression level as a driver of phenotypic penetrance and severity. Multiple copies of a cassette coding for pan-neuronal wildtype tau or T231E were inserted at a genomic safe harbor loci to create a repertoire of strains expressing tau from low to high levels. In stereotypical behavioral assays of locomotory activity, T231E selectively impacted phenotypic severity compared to wild-type human tau controls, which further tracked with age and tau expression level. However, deficits in associative memory were non-selective between tau and T231E. Moreover, genetic, pharmacologic, and molecular approaches indicated that mitophagy modulation could suppress T231E phenotypes. Additionally, a robust mitochondrial unfolded protein response (UPRmt) occurred in T231E, and loss of *atfs-1*, a transcription factor central to the UPRmt suppressed T231E toxicity. These results demonstrate that phenotypic severity is invariably associated with tau dosage, while early-AD relevant modifications can be causative drivers of selective deficits. Consistent with recent findings, enhancing mitophagy or suppressing potentially maladaptive consequences of persistent UPRmt induction can be beneficial. This provides a solid foundation for further interrogation into mitochondrial quality control disruption as a potential root cause for AD pathogenesis.

**Highlights:**

- Matched sets of pan-neuronal, multi-copy tau strains enhance experimental control
- Phosphomimetic tau elicits selective behavioral and neuronal dysfunction
- Phosphomimetic tau triggers a unique mitochondrial unfolded response
- Tau depletion and mitochondrial interventions rescue observed deficits

**Visual Abstract:** 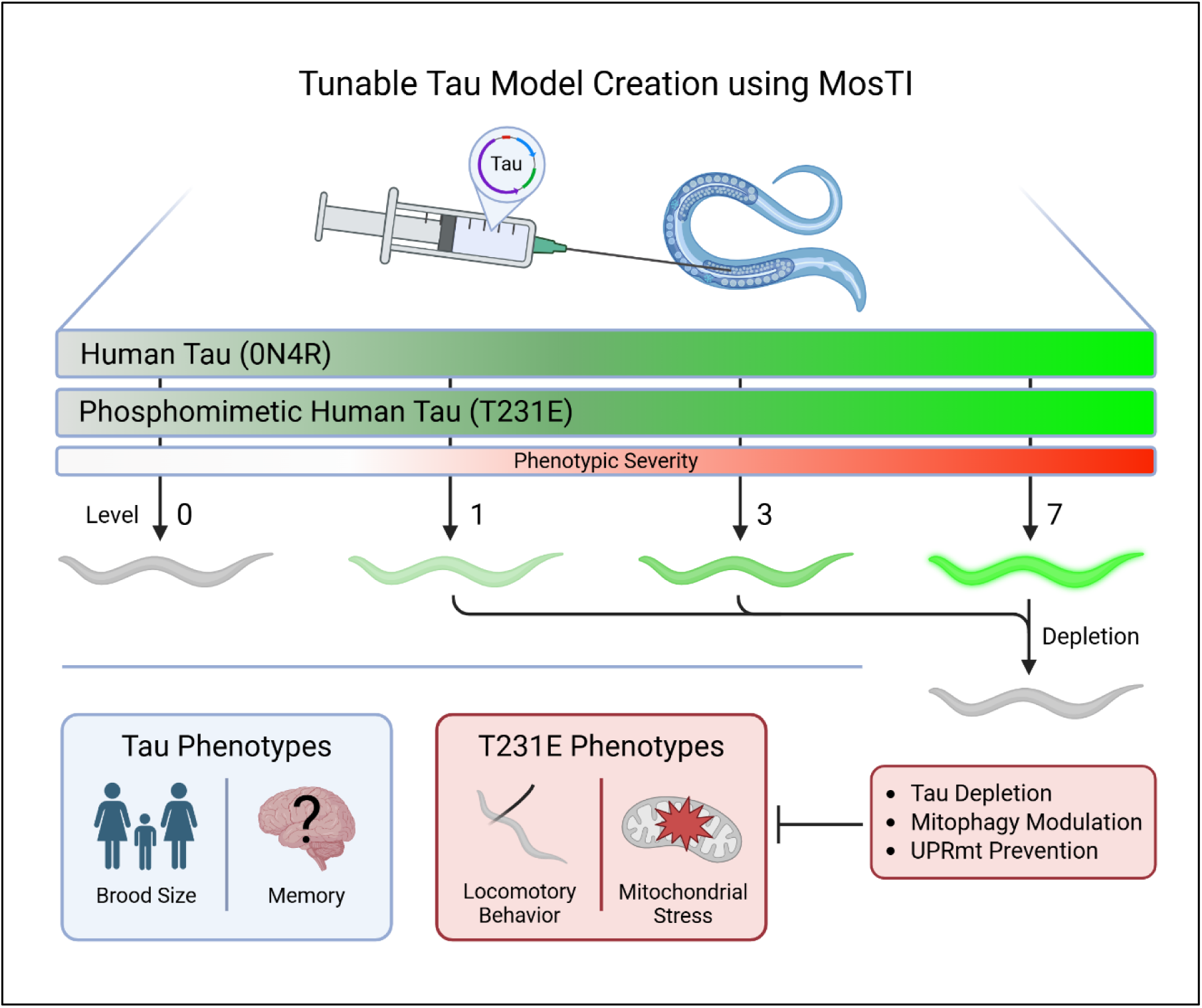

## Background and Introduction

Alzheimer’s disease is the most common form of dementia and presents a growing healthcare and economic burden worldwide ^1^. Hundreds of therapies are currently in clinical trials for Alzheimer’s disease (AD) ^2^, and approximately one third of current therapies are aimed toward treating AD symptoms rather than ameliorating underlying pathology. Among disease-modifying biologics, two thirds of registered therapies target some amyloid beta or tau isotype, hoping to stop AD progression at the root ^2^. Many of these biologics specifically seek to prevent aggregates from accumulating in patient’s brains. However, these trials depend on whether amyloid-beta or tau pathology are causative or merely correlative to AD progression, which is still poorly understood despite decades of research.

Many studies and models of choice have introduced aggregation-prone amyloid-beta and tau variants and shown how they impact neuronal homeostasis through central cellular processes such as autophagy, endocytic clearance, differentiation, synaptic transmission, cargo transport along microtubules, and mitochondrial function ^3^. However many of these models are fraught with caveats that may limit their translation to human disease: many mammalian models study familial AD mutations or tau isotypes associated with non-AD dementias, which limits their insight into AD pathology; cellular models fail to account for tissue wide pathologies, such as inflammation, that modify cellular dysfunction; and many studies in model organisms rely on overexpression to generate neurodegenerative phenotypes which may then be complicated by general proteostatic stress rather than protein-selective dysfunction.

Ample evidence now suggests that tau variants are critical and causative agents in human disease. Specifically, there are now three confirmed cases of human patients that possess familial AD mutations which are resistant to developing dementia ^4–6^. In each case, the patients’ brains possess aberrant amyloid-beta pathology but fewer tau neurofibrillary tangles than familial controls. These case studies highlight tau pathology as an essential contributor to cognitive impairment in AD, and recent studies have demonstrated that tau’s post-translational modifications (PTMs) are key modulators of its physicochemical properties and toxic potential ^7^. Thus, the many modifications which accumulate on tau before it manifests into aggregates may be critical events which trigger AD progression and warrant in-depth study.

Among early-disease PTMs, phosphorylation at T231 is of particular interest to AD pathology because it is currently being investigated as a clinical biomarker for AD diagnosis before symptoms manifest ^8, 9^. This is predominantly because it becomes enriched in brain regions that subsequently develop tau tangles, highlighting it as one of the earliest observed molecular changes in AD. Our lab has previously demonstrated that a phosphomimetic version of 0N4R human tau with threonine 231 mutated to Glutamic acid (T231E) causes neuronal dysfunction even when expressed at single-copy gene levels in *C. elegans* neurons ^10, 11^. Herein, to methodologically disentangle protein overexpression from selective toxicity of disease-associated tau modifications, we’ve generated a set of pan-neuronal, “tunable tau” *C. elegans* models where either wildtype or phosphomimetic T231E exhibit a wide range of expression levels depending on the strain and can be selectively depleted at will using an auxin inducible degron tag ^10, 12^.

We have leveraged this highly controlled set of multi-copy tau animals to address whether behavioral phenotypes exhibit selective vulnerability to the T231E phosphomimetic and to ask whether increased tau expression exacerbates these phenotypes in an age-dependent manner. We demonstrate that increased expression of T231E induces thrashing deficits not previously seen in single-copy models in a mimetic-specific manner, whereas wildtype tau and T231E are equally detrimental for an associative learning paradigm intended to mimic short term memory formation, with the magnitude of the deficit dependent solely on tau expression levels. We find that T231E-associated phenotypes can be suppressed by inducing mitophagy, indicating that T231E may selectively impact mitochondrial quality control. Moreover, high-level T231E expression drives a mitochondrial unfolded response (UPRmt). Perhaps most intriguing is our finding that genetic ablation of *atfs-1*, coding for a central transcription factor necessary for the UPRmt ^13^, fully suppresses T231E phenotypes. These studies suggest that chronic activation of the UPRmt selectively by a disease-associated mutant of tau can be maladaptive and provide a unique model to test whether boosting mitochondrial function may be a useful therapeutic avenue to combat AD.

## Materials and Methods

### C. elegans Growth and Maintenance

Nematodes were maintained at 20°C on Nematode Growth Media (NGM) plates and seeded with live *E. coli* strain OP50 which was cultured overnight at 37°C at 220 rpm, as previously described ^14, 15^. For experimental assays in which animals were age-synchronized, adults worms were treated with sodium hypochlorite to isolate embryos, and then 4th larval stage (L4) hermaphrodites were selected and moved to test plates. The day after moving was considered adult day 1. If assayed later in life, animals were transferred daily to avoid populations of mixed ages.

### Plasmid Construction

Briefly, transgenes from pTC10 (pUPN::AID::eGFP::*tbb-2* 3’UTR) and pTC11 (pUPN::AID::eGFP::TauT4::*tbb-2* 3’UTR) ^17^ were excised using Acc65I digestion and positively identified using SacI-HF digest. Transgenes were re-ligated into backbone vectors pTC10 and pTC11 ^17^, generating plasmids pTC14 and pTC15, respectively. Site-directed mutagenesis was performed to mutate threonine 231 in pTC15’s tau coding region to glutamic acid, thus generating pTC17 (pUPN::AID::eGFP::T231E::*tbb-2* 3’ UTR) (Quikchange II XL kit, Cat. # 200521, Agilent, Santa Clara, CA). Primers used for site-directed mutagenesis are listed in Table 2. Final plasmids were sequenced completely (Plasmidsaurus, San Francisco, CA) and used for transgene insertion, as described below.

### C. elegans Strain Generation Using MosTI Injection

For all strains herein (see Table 1), the wild-type background strain is Bristol-N2 and all lab-generated and externally acquired strains were outcrossed to Bristol-N2 at least four times. A spectrum of strains expressing various levels of multi-copy, pan-neuronal transgenes were created by integrating pTC14, pTC15, and pTC17 into strain CFJ42, a strain with a modified version of the Ch. II *ttTi5605* insertion site (*kstSi42*), using Mos-mediated Transgene Insertion (MosTI) ^16^. Briefly, transgenes from plasmids pTC14, pTC15, and pTC17 were amplified using traditional PCR using primers *ttTi5605* F2 and *ttTi5605* R2 (Table 2).

**Table 1.** *C. elegans* Strains Used in This Work.

| Strain Name | Genotype | Description/Purpose |
| --- | --- | --- |
| N2-Bristol | Wild-type | Control |
| KWN920 | *rnySi57 II; otIs730* | Single-copy, pan-neuronal AID-eGFP and pan-neuronal TIR1 |
| KWN921 | *rnySi58 II; otIs730* | Single-copy, pan-neuronal AID-eGFP-TauT4 and pan-neuronal TIR1 |
| KWN922 | *rnySi58(T231E) II; otIs730* | Single-copy, pan-neuronal AID-eGFP-T231E and pan-neuronal TIR1 |
| KWN987 | *rnyIs019 II: extrachromosomal array integration of pTC14[pUPN::AID::eGFP::tbb-2 3'UTR], pSEM319, and pCFJ782 into Chr. II* | Integrated via MosTI<sup>16</sup>; multi-copy, pan-neuronal eGFP; "G15" |
| KWN989 | *rnyIs026 II: extrachromosomal array integration of pTC15[pUPN::AID::eGFP::TauT4::tbb-2 3'UTR], pSEM319, and pCFJ782 into Chr. II* | Integrated via MosTI<sup>16</sup>; multi-copy, pan-neuronal TauT4; "T3" |
| KWN991 | *rnyIs028 II: extrachromosomal array integration of pTC15[pUPN::AID::eGFP::TauT4::tbb-2 3'UTR], pSEM319, and pCFJ782 into Chr. II* | Integrated via MosTI<sup>16</sup>; multi-copy, pan-neuronal TauT4; "T7" |
| KWN994 | *rnyIs038 II: extrachromosomal array integration of pTC17[pUPN::AID::eGFP::T231E::tbb-2 3'UTR], pSEM319, and pCFJ782 into Chr. II* | Integrated via MosTI<sup>16</sup>; multi-copy, pan-neuronal T231E; "E3" |
| KWN996 | *rnyIs040 II: extrachromosomal array integration of pTC17[pUPN::AID::eGFP::T231E::tbb-2 3'UTR], pSEM319, and pCFJ782 into Chr. II* | Integrated via MosTI<sup>16</sup>; multi-copy, pan-neuronal T231E; "E7" |
| KWN1021 | *rnyIs019 II; otIs730* | G15 x TIR1 |
| KWN1022 | *rnyIs026 II; otIs730* | T3 x TIR1 |
| KWN1023 | *rnyIs028 II; otIs730* | T7 x TIR1 |
| KWN1024 | *rnyIs038 II; otIs730* | E3 x TIR1 |
| KWN1025 | *rnyIs040 II; otIs730* | E7 x TIR1 |
| KWN1031 | *otIs730 [pUPN::TIR1::mTur2 + inx-6(prom18)::tagRFP]* | Pan-neuronal TIR1 for AID depletion (outcrossed to N2 from strain OH15845) |
| OH15495 | *otIs696* (NeuroPAL) | Used for polychromatic identification of all neurons |
| KWN1035 | *otIs696; rnyIs040 II; otIs730* | E7 x NeuroPAL |
| CL2355 | *tm376* (912 bp promoter deletion) | *dct-1* loss-of-function (lf) |
| KWN1038 | *rnyIs040 II; otIs730; tm376* | E7 x *dct-1* (lf) |
| AU529 | *tm4525* (432 bp deletion + 7 bp insertion) | *atfs-1* loss-of-function (lf) |
| KWN1042 | *rnyIs040 II; otIs730; tm4525* | E7 x *atfs-1* (lf) |

**Table 2.**
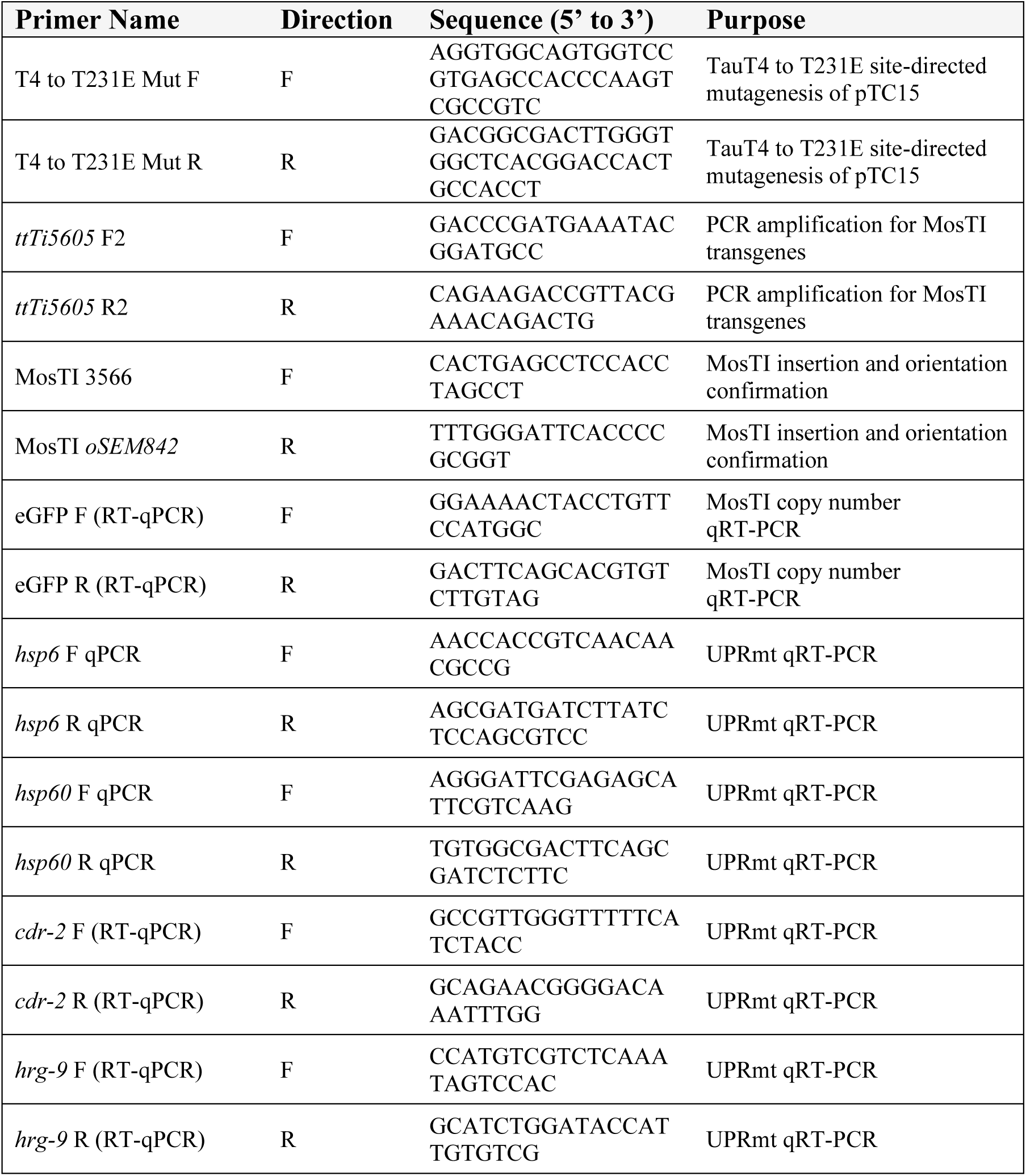
List of Primers Used in this Study.

The PCR products were micro-injected into the germline alongside targeting vector pSEM319 (MosTI targeting vector with a split-unc-119 rescue gene fragment, Addgene #159823) and selection marker pCFJ782 (Hygromycin resistance gene, Addgene #190933). Injected animals were then moved to individual plates and the resulting extrachromosomal array strains were maintained on NGM plates containing 0.5 mg/mL Hygromycin B (Cat. # 10687-010, Invitrogen, Carlsbad, CA) for two weeks.

In a second round of injections, CRISPR-Cas9 gene editing was used to integrate the extrachromosomal arrays into Chr. II site *kstSi42* via non-homologous end joining (NHEJ) by cutting the insertion site and target vector within the array. Targeting crRNAs were from Horizon Discovery (Cambridge, UK) and were complexed to scaffolding RNAs for Cas9, with the following recognition site:

*cbr-unc-119* MosTI targeting RNA, 5’-GATATCAAGAGTTAGTTGAG-3’

The resulting strains were selected via split-*unc-119* rescue and pan-neuronal eGFP fluorescence and homozygosed for the insertion. In total, at least eight strains were generated for each of the three targeted insertions (eGFP, TauT4 and T231E). Five of these were selected based upon matched protein expression levels and selected for further analysis. These strains were used to confirm successful gene insertion in the correct orientation via genomic PCR with primers MosTI 3566-F and MosTI *oSEM842*-R (Table 2) and backcrossed 4x to N2 prior to further use.

### Fluorescent Intensity Quantification

Day 1 adult animals were imaged using a Nikon Eclipse inverted microscope coupled to a six channel LED light source (Intelligent Imaging Innovation, Denver, CO), an ORCA-Flash4.0 V2 Digital CMOS camera (Hamamatsu Photonics, Bridgewater Township, NJ) and Slidebook6 software (Intelligent Imaging Innovation, Denver, CO). All images were acquired under the same exposure conditions, and each experiment was imaged in one session. Fluorescent intensities were quantified by creating a mask encompassing the entirety of the fluorescence surrounding the animal’s nerve ring and measuring the background subtracted mean intensity.

### Neuronal ID Using NeuroPAL

Day 1 adults were mounted on 2% agarose pads on glass slides and immobilized with 1 mM tetramisole hydrochloride (#L-9756; Sigma-Aldrich, St. Louis, MO). Imaging was performed using a Laser Scanning Confocal microscope (Inverted Leica DMi8 and LAX software). All images were acquired using a 20x/0.75 objective (Air – HC PL Apochromat C52). Briefly, tail neurons were imaged, image volumes were constructed using Imaris Viewer, and neurons were manually identified using NeuroPAL markers, as described ^18^.

### Lifespan Analysis

Animals were synchronized via bleach spot and maintained on regular NGM plates until day 1 of adulthood. Then, approximately 30 animals per strain per replicate were transferred to plates containing 50 μM 2’-deoxy-5-fluorouridine (FUdR) to suppress fertility and then moved to fresh FUdR plates every three days to ensure healthy living conditions. Animals were counted each day until none were left alive.

### AID Depletion

Animals were transferred to either regular NGM plates or NGM plates containing 4 mM indole-3-acetic acid (IAA, A10556; Thermo Scientific Chemicals, Haverhill, MA) and left to deplete for 24 hours before assaying, as described previously ^17^.

### Touch Sensitivity Assay

The behavioral response to being touched by an eyelash was adapted from an assay previously described ^19^. The animals were touched anteriorly specifically behind the terminal bulb of the pharynx with an eyelash 10 times per animal, with a 10 s gap between each touch. Typically, if the animal demonstrates an omega turn or if it reversed its direction after an anterior touch, the animal was scored as giving a positive response. Touch response percentage was generated by the number of times an animal responded to the touch stimulus over the total number of times they were touched. All touch response assays reported here were blinded by an independent researcher before experimentation.

### Thrashing Assays

A drop of 2% agarose (BP164-500; Thermo Fisher Scientific) was poured over a glass slide and allowed to dry and then 20 μl of M9 was poured on it. Age-synchronized animals were picked to that drop of M9 buffer. After 2 min in M9, thrashing rates were counted and recorded manually. A single thrash was defined as a complete change in the direction of the body down the midline. Animals that were motionless for 10 s were discarded from the analysis. For assays using paraquat (PQT), animals were transferred to NGM plates containing 8 mM PQT the day before the assay and treated overnight. In cases where animals were treated with celastrol, animals were moved to NGM plates containing either 0.025% DMSO (vehicle) or 25 µM celastrol (70950, Cayman Chemical Company, Ann Arbor, MI) in 0.025% DMSO the day before assaying and treated overnight.

### Associative Memory Assays

Briefly, for each experimental strain, day 1 adults were washed in M9 and transferred to conditioning NGM-agar plates (without food) in the presence or absence of isoamyl alcohol (IA, AX1440-1, EM Science, Gibbstown, NJ) and left to starve for 90 minutes. Conditioned animals were then washed with M9 and transferred to assay plates to assess their chemotactic attraction or aversion to isoamyl alcohol, as previously described ^20^. The chemotactic index is calculated as (IA -T)/ (IA + T + S), where IA equals the number of worms residing in the quadrant containing IA, T equals the number of worms residing in the quadrant opposite the IA quadrant, and S equals the number of worms within the starting quadrant, at the end of the assay period.

### Brood Size Assays

For each replicate, three L4 larvae were moved to a new, small NGM-plate each day until the animals stopped laying embryos (typically by day 5 or day 6 of adulthood). Embryos were counted each day just after transfer, and the reported brood sizes for each replicate are the summation of all embryos laid by the three animals throughout the entire brood-bearing window. For transgenerational depletion studies, F0 animals were transferred as embryos to 4 mM IAA NGM plates to deplete T231E tau throughout their life. For each subsequent generation, the larvae laid at day 1 of adulthood for the previous generation were maintained on IAA until adulthood and used for brood size assays.

### RNA Extraction and qRT-PCR

For each sample, approximately 2,000 - 3,000 animals at day 1 of adulthood were collected in M9, washed 3x, and the supernatant was discarded after letting the animals settle on ice. 1 mL Trizol® Reagent (15596026, Ambion, Austin, TX) was added to extract RNA, animals were flash frozen in liquid N2 and thawed in a 37°C water bath four times. Samples were moved to ice, then 200 µL chloroform (C2432, Sigma Aldrich, Burlington, MA) was added and samples were shaken vigorously and centrifuged at 12,000 rcf for 15 minutes at 4°C. The supernatant was discarded, 500 µL IPA was added, and the samples were incubated at room temperature for 10 minutes and centrifuged at 12,000 RCF for 15 minutes at 4°C. The supernatant was discarded, 1mL ice-cold 75% EtOH was added, the samples were vortexed for 30s, and then the samples were centrifuged at 7,500 rcf for 5 minutes at 4°C. The supernatant was removed and all EtOH was allowed to evaporate before reconstituting the RNA pellet in 50 µL RNA-ase free water. Contaminating genomic DNA was digested using a Turbo DNA-free™ Kit (AM1907, Invitrogen, Carlsbad, CA), the RNA concentration of each sample was measured via NanoDrop (LT1022, Thermo Scientific, Waltham, MA). 300ng RNA was used to create cDNA libraries from each sample using Quantabio qScript cDNA Supermix (95048, Quantabio, Beverly, MA). qRT-PCR was run in 96-well plates using 2x Universal SYBR Green Fast qPCR Mix (RK21203, ABclonal, Woburn, MA) and the primers listed in Table 2.

### Statistical Analysis

All statistical analyses were conducted using Prism 8.0 (GraphPad Software Inc, Boston, MA), with alpha-error level of p < 0.05 considered to be significant. Data were averaged and represented as mean ± standard error (mean ± SEM) or as mean ± standard deviation (mean ± SD), depending upon the number of experimental replicates. In general, differences between matched pairs of samples were analyzed using student t-tests, and group differences were analyzed with either one-way or two-way ANOVA, depending upon the variables. For all tests herein, *P < 0.05, **P < 0.01, ***P < 0.001, ****P < 0.0001.

## Results

### Tunable Tau *C. elegans* Models Possess Selective Deficits in High-Copy T231E Expressing Animals

Historically, most *C. elegans* models of protein aggregate neurodegeneration have relied on overexpression of exogenous proteins, with robust expression levels driving visible anatomical and behavioral phenotypes. Here, we adopted a different approach, creating a repertoire of strains that express tau at varying levels. Our expectation was that selective vulnerability in terms of neuronal dysfunction or heightened sensitivity to pathologic AD-associated tau mutants would be more readily assessed across a dosage spectrum. Since tau is prone to aggregation and can create neuropathology if expressed at a high enough level, even when it is unmodified, we further anticipated that this model would be useful for studying incipient tau pathology prior to aggregation.

Previously, we showed that strains which express single-copy levels of human tau under control of the Ultra Pan-Neuronal (UPN) promoter ^18^ exhibited phenotypes specific to a T231E phosphomimetic ^17^. However, expression under the UPN promoter is weak, leading us to wonder whether methodologic increases in expression might unveil phenotypes previously described in other pan-neuronal tau models ^21–23^ and whether those phenotypes would also be selective for T231E. Hence, we used a technique called Mos-mediated Transgene Insertion (MosTI) ^16^ to create strains expressing fusions of an auxin inducible degron (AID) to eGFP, eGFP::TauT4, or eGPF::T231E (**Fig. 1A**). Integration through MosT1 occurs via NHEJ mediated recombination from CRISPR/Cas9 cleavage fragments of an extrachromosomal array. The size of the inserted DNA fragment ranges greatly depending upon the frequency and stochastic distribution of CRISPR target site in the array compared to the transgenic cassette expressing the gene of interest.

**Figure 1.**
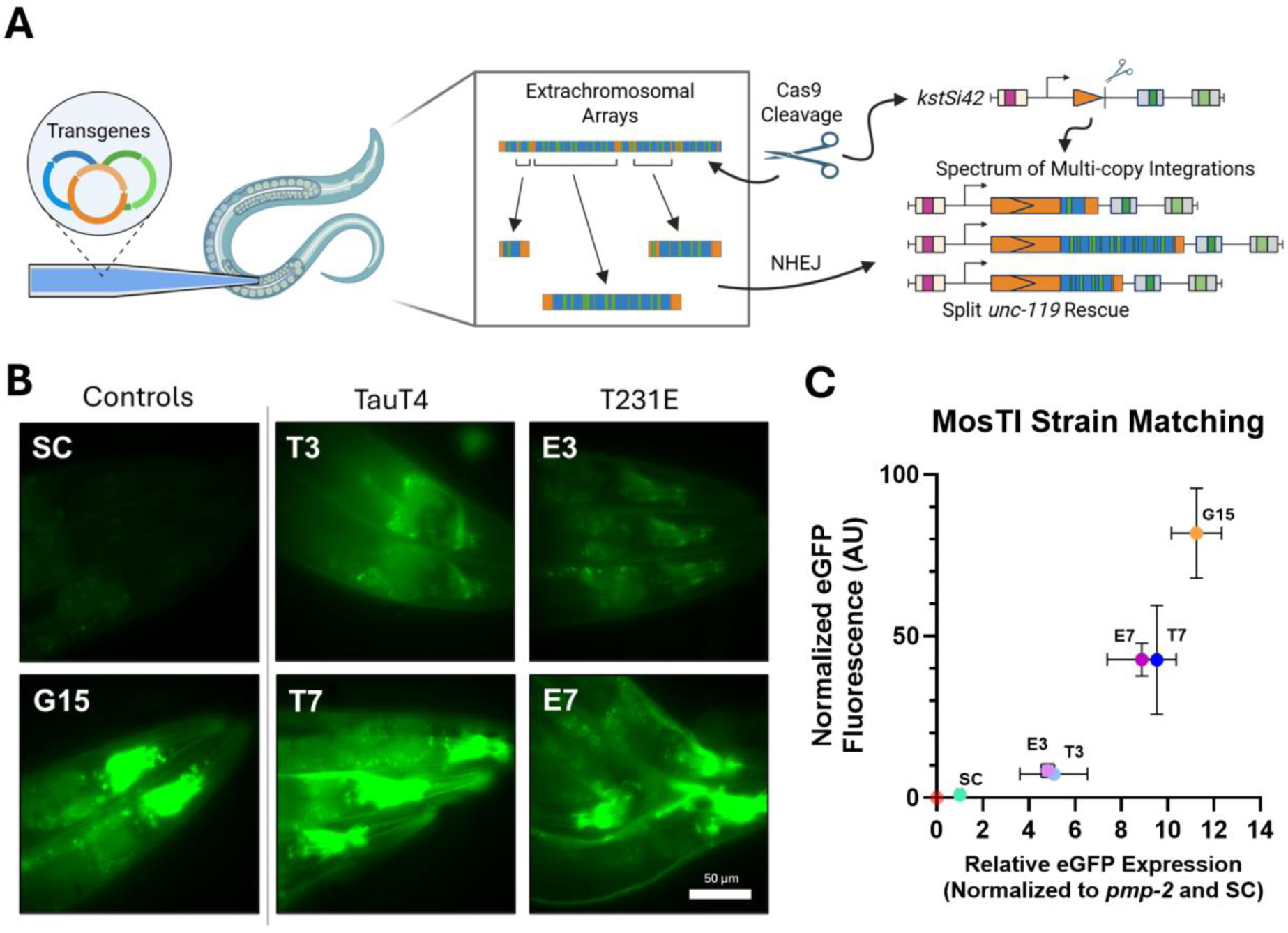
Creation of Tunable Tau Model Via MosTI Injection. (A) Schematic summarizing the MosTI injection procedure and how it leads to the generation of strains with a spectrum of integrated transgene copy numbers into safe-harbor loci *kstSi42*. (B) Fluorescent images acquired near the nerve ring of transgenic animals as labeled demonstrating the differences in intensity among the low and high-copy strains. All images are of eGFP fluorescence, either alone or fused to tau. Images were taken on the same day, under the same exposure condition, and scaled identically for presentation. SC is single copy T231E; G15 is very high level eGFP alone; T3 and E3 are medium-level TauT4 and T231E, respectively; T7 and E7 are high-level TauT4 and T231E. Scale bar represents 50 μm. (C) Quantification of eGFP fluorescent intensity (y-axis) and eGFP expression using qRT-PCR (x-axis) to match MosTI strains. Mean fluorescent intensities were normalized to SC intensity values and qRT-PCR values were normalized to housekeeping gene *pmp-2* and then to *pmp-2* normalized SC transcript eGFP expression. For intensity measurements, N=3-5 animals. For qRT-PCR, N=3 technical replicates.

In practical terms, this means that it is possible to recover inserts that range from near single-copy (SC) to many copies. It is important to note that the MosT1 derived strains express less target protein than the starting extrachromosomal arrays themselves, making this a perfect model to avoid the confounds of non-physiologic overexpression.

We recovered at least eight independently derived strains for each target gene ranging in fluorescent intensity from low to high, with the spectrum of strains expressing GFP generally brighter than those expressing TauT4 or T231E. For the purposes of this work, we focused on four strains with matched expression levels for TauT4 and T231E (medium and high) to facilitate their comparison and chose one bright GFP strain to serve as a control (**Fig. 1B**). These strains were colloquially named using an “XY” convention, where X = (G, T, or E, standing for eGFP, TauT4, or T231E, respectively) and Y = an estimated relative fluorescent intensity score. A plot of quantified fluorescent intensity versus normalized relative mRNA suggests that the paired TauT4 and T231E strains (T3 and E3; T7 and E7) expressed nearly equal amounts of both transcript and protein, while the eGFP strain expressed significantly more than even the highest tau strains (**Fig. 1C**).

We previously showed that animals expressing single-copy levels of T231E, but not TauT4, selectively exhibited age-dependent deficits in the locomotory response to gentle touch but did not exhibit deficits in associative memory or thrashing behavior ^17^. Associative memory in *C. elegans* can be assayed through chemotactic learning, where worms learn to associate a chemical cue with food or starvation, then show altered attraction or avoidance in a chemotaxis assay ^24^. For example, N2 animals normally attracted to isoamyl alcohol (IA) exhibited robust negative chemotaxis immediately following preconditioning with IA during starvation (**Fig. 2A**).

**Figure 2.**
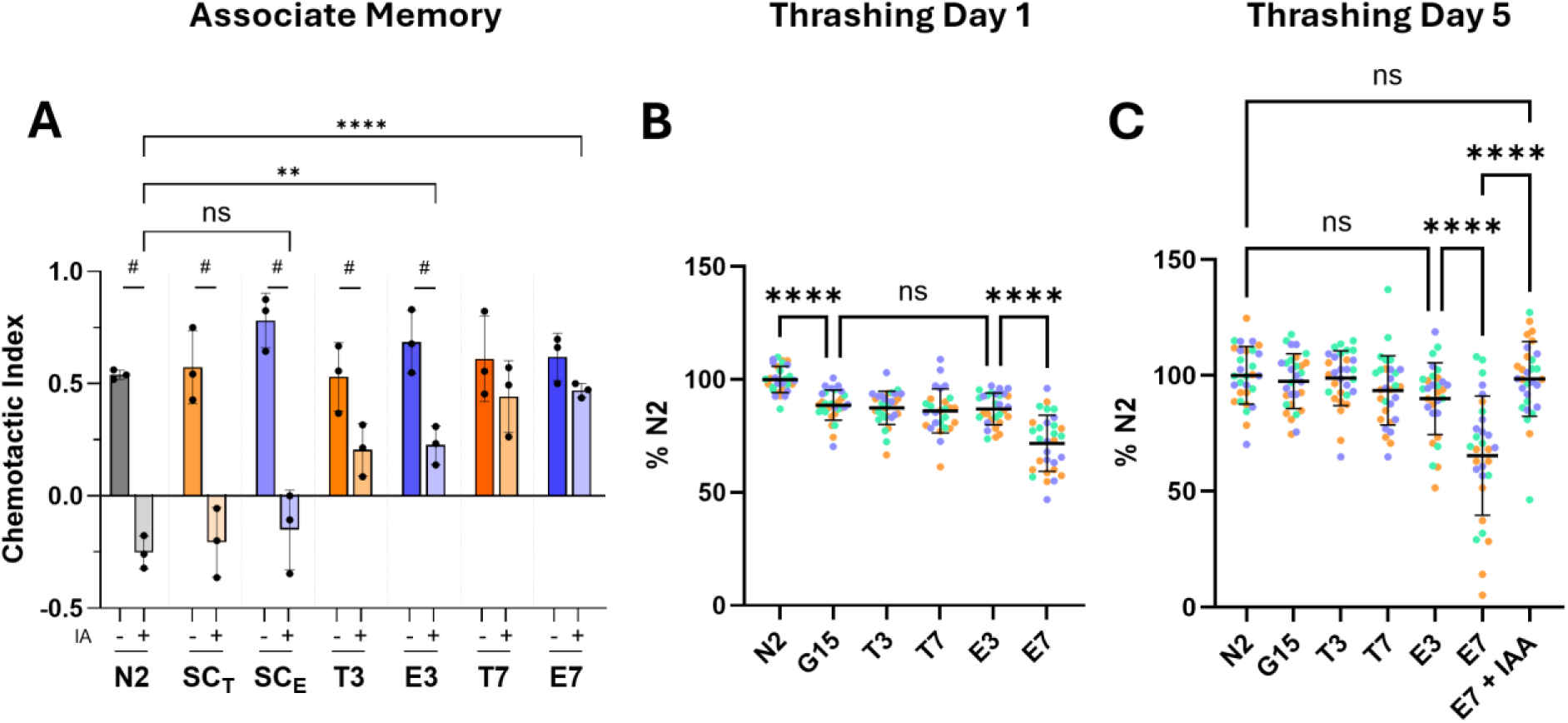
Selectivity of Phenotypic Responses in Tunable Tau Strains. (A) Associative memory assay. Isoamyl alcohol (IA) is normally an attractant for *C. elegans*. Chemotactic indexes were calculated for IA in animals preconditioned by starvation in the absence (“-”) or presence (“+”) of IA. “#” denotes a significant change in chemotactic index between paired “-” and “+” animals, reflecting short-term associative memory of starvation causing an agnostic or repulsive response to IA. Each point represents a population of 50-100 animals, N=3 biological replicates. Individual strains with or without conditioning were compared using unpaired students t-tests. Chemotactic indexes between strains, yet within the same treatment group, were compared using a one-way ANOVA followed by Tukey-corrected multiple comparisons. F = 18.24, DF = 20. (B and C) Thrashing assays. Rates are shown for day 1 (panel B) and day 5 (panel C) animals as a percentage of the average N2 control rate. To test reversibility, E7 animals were treated overnight at day 4 with 4mM IAA to trigger auxin-inducible depletion and assayed at day 5. The plots show mean ± SD, with each point representing an individual animal. N = 30 animals from three biological replicates. Each biological replicate is denoted by a unique color. Statistical analysis was performed using a one-way ANOVA followed by Tukey-corrected multiple comparisons. F = 29.05 (B) and 15.70 (C). DF = 206 (B) and 289 (C).

As reported previously ^10^, single copy strains T1 and E1 were indistinguishable from N2 and displayed a normal associative response to preconditioning with starvation and IA together (**Fig. 2A**). However, strains T3 and E3 exhibited a moderate deficit in associative memory formation while strains T7 and E7 were both statistically unresponsive to preconditioning (**Fig. 2A**). These results clearly demonstrate the impact of tau dosage on phenotypic severity. Moreover, TauT4 and T231E performed similarly in this assay when paired by expression level, suggesting that unlike previously described phenotypes, this phenotype was not selective for T231E. We note that none of the strains, including those with associative memory deficits, exhibited a defect in primary chemotaxis to IA in the absence of associative preconditioning, suggesting that odorant detection and the ability to locomote appropriately were unaffected (**Fig. 2A**).

We also found deficits in thrashing behavior that were not observed in SC strains. In this case, the defects were restricted to animals expressing the highest level of T231E, with E7 animals exhibiting significantly lower thrashing rates compared to all other strains at days 1 and 5 (**Fig. 2B and 2C**). Of note, simply expressing eGFP pan-neuronally at extremely high levels caused minor touch and thrashing deficits at day 1; however, these differences were not significant at day 5 (**Fig. 2B and 2C**). In order to demonstrate that the E7 defect was specific to tau expression, we show that AID depletion of T231E restored thrashing to N2 levels at day 5 (**Fig. 2C**). Finally, we assessed touch response in the multicopy strains and found a similar phenotypic selectivity for T231E (**Fig. S1**). Both of these findings mirror the results of similar approaches published using SC strains ^17^.

Additionally, we noted that T7 and E7 strains had a markedly reduced brood size. Hermaphroditic *C. elegans* self-mate to generate ∼300 progeny per animal, while the T7 and E7 strains produced roughly half or a third of that, respectively (**Fig. S2**). The brood size defect was limited to the highest-level multi-copy strains (data not shown). Tau depletion did not rescue the brood size deficit in the first generation (**Fig. S2)**, however, we continued to assess brood size over the next four generations, with embryos from mothers raised on auxin being laid onto auxin-containing plates so that the worms never experienced tau accumulation over multiple generations. We were astounded to find that partial rescue of the brood size phenotype was observed in the fourth generation and complete rescue was observed one generation later (**Fig. S2**), which is consistent with *C. elegans* exhibiting transgenerational stress responses that persist for a similar period ^25–28^. Clearly, the relevance of these observations to AD is uncertain, since it is an age-dependent, post-reproductive disease. However, the formal possibility exists that this may represent a cell-nonautonomous stress response to pathologic tau accumulation in neurons.

Similarly, there remains a formal possibility that behavioral phenotypes could be mediated via neuronal loss, either through a lack of neuron formation during development or through neuronal cell death, albeit unlikely due to the ability to recover function through AID depletion of tau. To assess the presence or absence of neurons, we used a multicolor transgene called neuronal polychromatic atlas of landmarks, or “NeuroPAL” ^18^. NeuroPAL expresses four fluorophores (mTAGBFP2, CyOFP1, mNeptune2.5, and TagRFP-T) under a gamut of neuronal promoters, ultimately creating a pan-neuronal identification scheme which allows researchers to visually distinguish individual neurons within dense ganglia using a combination of neuron position and unique neuron color. Of note, the fluorophores within NeuroPAL are fluorescently distinguishable from eGFP, allowing for simultaneous imaging of our eGFP tagged tau. We crossed our high-copy T231E phosphomimetic transgene E7 into the NeuroPAL background (OH15495) and imaged the tails of several day 1 animals to determine if any neurons were missing when behavioral deficits began to manifest. We found no evidence of missing neurons in day 1 E7 animals (**Fig. 3, Supplemental Table 1, and Supplemental Videos 1 and 2**). These results confirm that T231E does not drive appreciable neuronal cell loss in the E7 strain.

**Figure 3.**
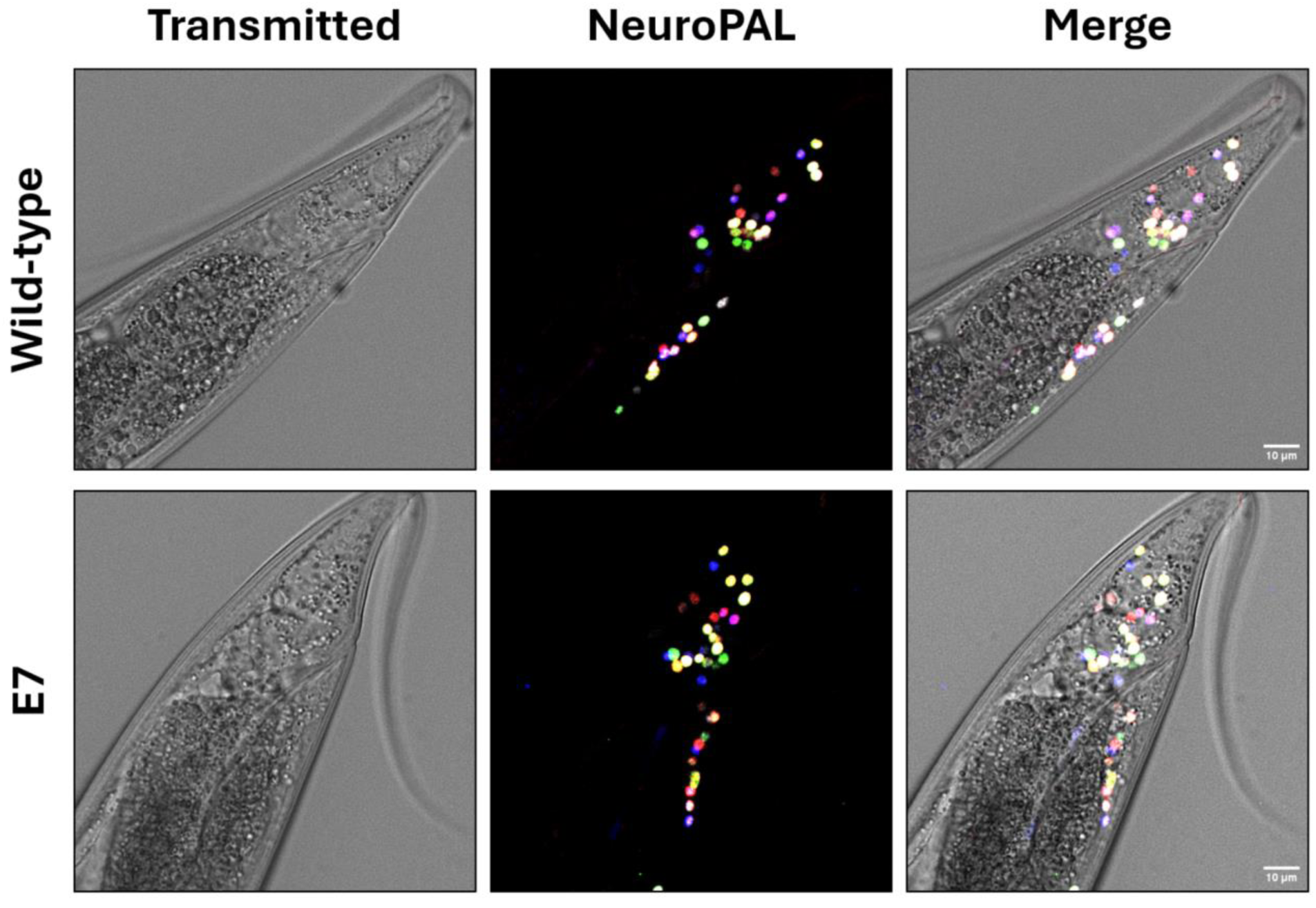
T231E-mimetic Animals Lack Overt Neurodegeneration. Representative images of neurons present in wild-type (OH15495) and high-copy phosphomimetic “E7” animals (KWN1035) at day 1 of adulthood. Neurons were identified using NeuroPAL, a “Polychromatic Atlas of Landmarks”, designed to identify all neurons within *C. elegans* using a combination of neuron position and unique fluorescent signatures using four neuronally expressed fluorophores, as described ^18^. Three-dimensional rotating images of these strains, including tau fluorescence, are shown in Supplemental Video 1. Quantitative data with individual neuronal identification / presence are presented in Supplemental Table 1. Scale bars represent 10 µm.

### High-Copy Expression of T231E Uniquely Impacts Mitochondrial Quality Control and Stress Signaling

Others have reported previously that enhancing mitophagy can rescue neurological deficits in AD models ^24, 29^, and we have previously shown that even single-copy expression of T231E is sufficient to suppress mitophagy in neurons ^30^. Thus, we next wondered whether inducing mitophagy could also rescue the touch and thrashing deficits caused by T231E expression. We induced mitophagy by treating with celastrol, a potent mitophagy inducer currently being investigated in clinical trials, ^31^ and found that it completely rescued thrashing and touch deficits at day 1 (**Fig. 4A and 4B**). Since restoring mitophagy restored touch and thrashing behavior back to wild-type levels, we hypothesized that T231E may be selectively impacting *dct-1*, a key regulator of mitophagy and the *C. elegans* ortholog of NIX/BNIP3L ^32^. We crossed E7 animals and G15 controls into *dct-1* loss of function animals and found that *dct-1* knockout completely rescued T231E induced touch deficits at day 1 (**Fig. 4D**) and rescued thrashing deficits at day 1, albeit partially (**Fig. 4C**). Of note, *dct-1* knockout also mildly increased thrashing rates of N2 animals (**Fig. 4C**). Finally, we wondered whether E7 animals may be more susceptible to mitochondrial stress and treated them with 8 mM PQT to induce redox cycling and ROS production, as described previously ^33^. Surprisingly, we found that wild-type N2 animals were susceptible to the mitochondrial stress imparted by PQT, yet the E7 animals seemed resistant, as indicated by thrashing analysis at day 1 (**Fig. 4E**).

**Figure 4.**
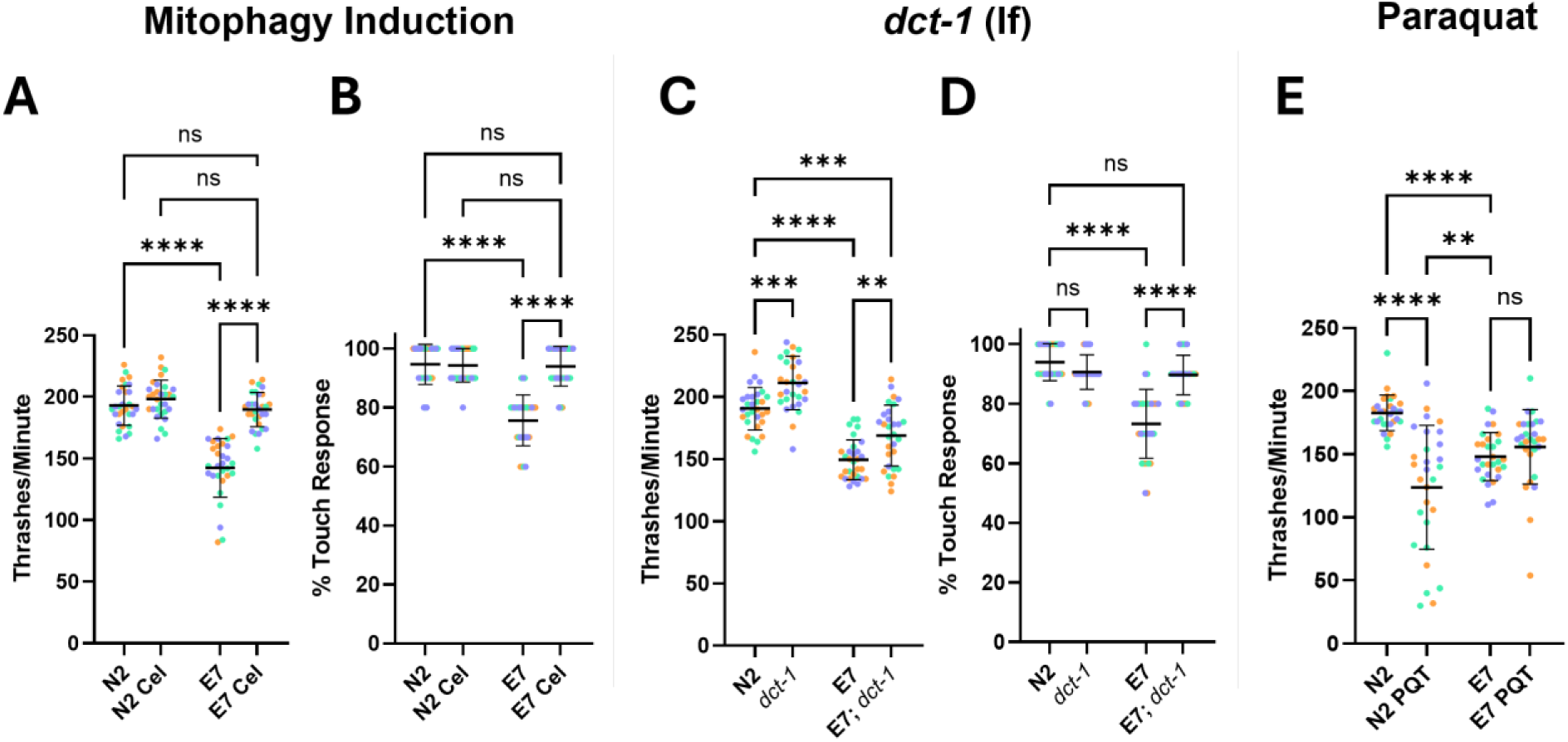
Mitophagy Impacts the Phenotypic Severity of T231E. (A) Quantification of thrashing rates and (B) touch responsiveness in day 1 adult N2 control and E7 (AID::eGFP::T231E) animals after treatment with vehicle (DMSO) or celastrol overnight. (C) Quantification of thrashing rates and (D) touch responsiveness in day 1 adult N2 control and E7 (AID::eGFP::T231E) animals as a function of the mitophagy receptor gene *dct-1*. (E) Quantification of thrashing rates in day 1 adult N2 control and E7 (AID::eGFP::T231E) animals after treatment with 8mM PQT overnight. Each point represents an individual animal, N = 30 animals from three biological replicates. Each biological replicate is denoted by a unique color. Statistical analysis was performed using a one-way ANOVA followed by Tukey-corrected multiple comparisons. F = 64.47, 52.86, 53.15, 40.90, 17.81 respectively. DF = 119 (A-E).

Cellular stress responses often involve the induction of compensatory signaling pathways that help restore homeostasis and confer resistance to subsequent similar stressors. Given that mitochondrial stress is a ubiquitous facet of neurodegenerative diseases including AD, we hypothesized that resistance to PQT stress might occur through T231E triggering adaptive responses surrounding mitochondrial fitness. The mitochondrial unfolded protein response (UPRmt) acts to restore mitochondrial health, and in vertebrates plays a critical role in metabolism, immunity, and diseases like cancer, diabetes, and neurodegeneration ^34–36^. *C. elegans* also possess a robust UPRmt that has been shown to impact stress resistance, innate immunity, and lifespan, among other phenotypes ^37, 38^. qRT-PCR was used to measure transcript levels for several genes which have known increases in expression in response to stimulation of the UPRmt (**Fig. 5A, 5B, 5C, and 5D**). Our results clearly demonstrate that the canonical UPRmt markers *hsp-6* and *hsp-60* were elevated in both T7 and E7 strains (**Fig. 5A and 5B**). However, other genes recently shown to be induced through UPRmt signaling ^37, 38^ exhibited different profiles (**Fig. 5C and 5D**). For example, *cdr-2*, a cadmium responsive protein involved in heavy metal detoxification, was unresponsive to either T7 or E7 (**Fig. 5C**) whereas *hrg-9*, a haem chaperone which is implicated in other mitochondrial disorders ^39^, is uniquely upregulated in E7 animals (**Fig. 5D**). Notably, qRT-PCR was performed on whole animals, thus we cannot differentiate between a cell non-autonomous responses across the entire animal or a stronger responses occurring selectively in the neurons.

**Figure 5.**
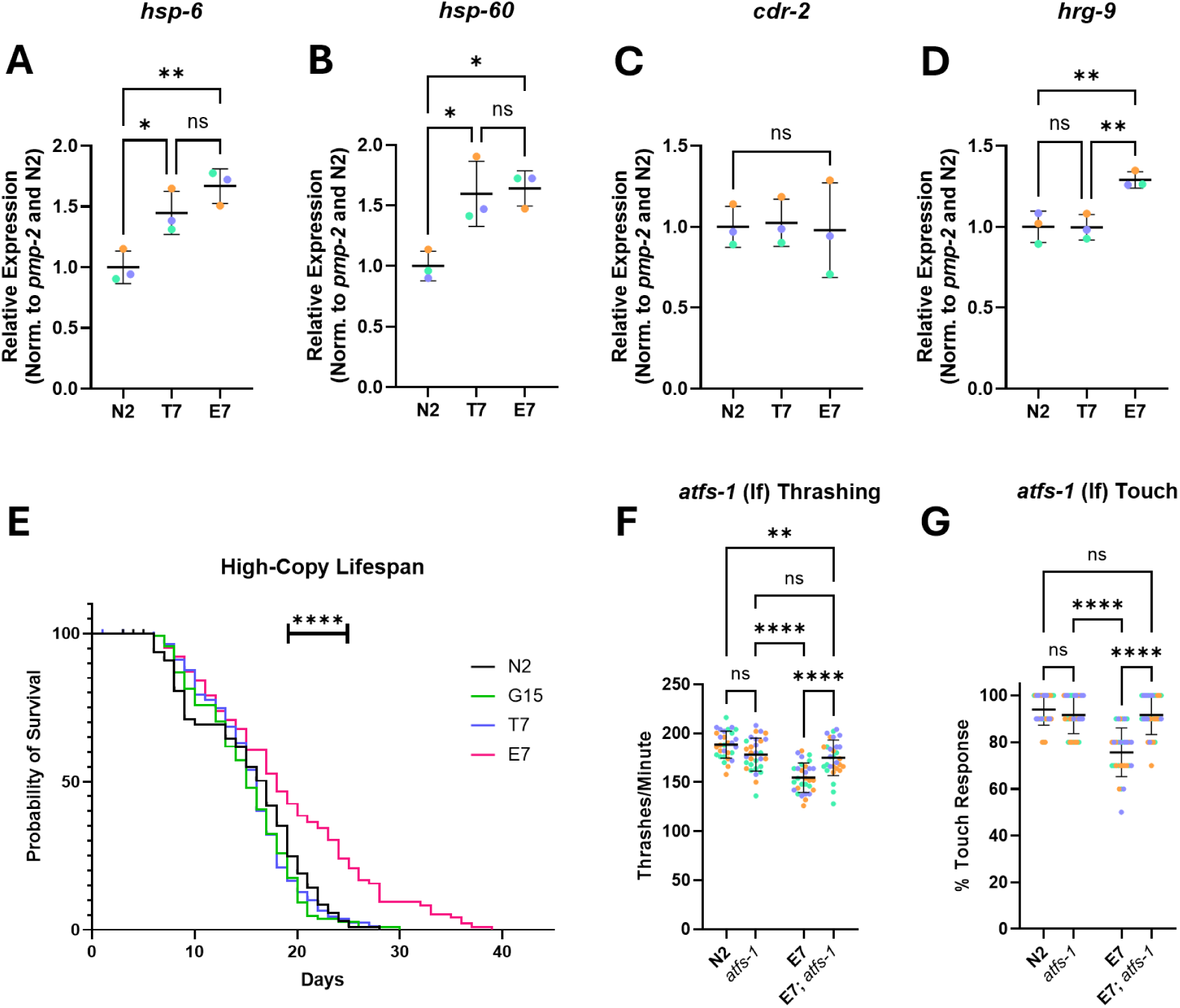
High-Copy T231E Induces Maladaptive UPRmt Signaling. (A-D) Quantification of RNA abundance for UPRmt genes in day 1 animals via qRT-PCR, normalized to housekeeping gene *pmp-2* and then N2 ratios. Primer pairs for each target gene are listed in Table 2. N = 3 biological replicates of approximately 2,000-3,000 animals. Each biological replicate is denoted by a unique color. Statistical analysis was performed using a one-way ANOVA followed by Tukey-corrected multiple comparisons. F = 14.99, 10.65, 0.04, and 14.09, respectively. DF = 8 (A-D). (E) Kaplan-Meier survival curve for strains N2, G15, T7 and E7 animals. Curves were calculated using 4 biological replicates of 25-30 animals each. Median lifespan for N2, G15, T7, and E7 were 17, 15, 16, and 18 days, respectively. (F) thrashing rates and (G) touch response data in day 1 adult N2 control and E7 (AID::eGFP::T231E) animals as a function of the central UPRmt transcription factor gene *atfs-1*. N = 3 biological replicates of approximately 10 animals. Each biological replicate is denoted by a unique color. Statistical analysis was performed using a one-way ANOVA followed by Tukey-corrected multiple comparisons. F = 23.09 (F) and 30.05 (G). DF = 119 (F-G).

Since mitochondrial stresses, including dysfunctional electron transport, ROS generation, hypoxia, and nutrient deprivation, elicit adaptive responses that can extend *C. elegans* lifespan ^40^, we tested whether T7 and E7 worms lived longer than G15 or N2 controls (**Fig. 5E**). A significant extension of lifespan was only observed in E7, but not T7 populations (**Fig. 5E**). Interestingly, the trace for E7 animal lifespan differs mainly after 20 days of age, indicating that animals which make it to advanced age specifically live longer, whereas mortality rates are similar up to that point.

To further examine the impact of UPRmt activation on tau related phenotypes, we leveraged a deletion allele of *atfs-1*, which normally codes for a transcription factor that is central to induction of the UPRmt in *C. elegans* ^13^. We found that the loss of *atfs-1* suppressed both the E7 thrashing (**Fig. 5F)** and touch response deficits **(Fig. 5G**), consistent with the idea that normally protective signaling pathways can become maladaptive through chronic activation.

## Discussion

*C. elegans* models of neurodegenerative disease often rely on expressing sufficient amounts of a disease relevant protein to elicit an observable phenotype ^41, 42^. Many advances have been made possible via this approach due to *C. elegans*’ ease of use, its well-characterized nervous system, and its conserved machinery surrounding critical neuronal processes, such as mitochondrial quality control, as well as the much lauded power of worm genetics ^21, 22, 37, 43, 44^. Though expressing tau in *C. elegans* neurons is convenient approach to assessing its role in tauopathies, conclusions drawn from this approach require careful interpretation, as what is deemed as appropriate protein expression levels may be defined retrospectively - by the ability to trigger a measurable phenotype - rather than by physiological relevance. Thus, it is almost impossible to determine the “appropriate” level of expression, particularly in a model organism that does not natively express these proteins. These limitations should be considered when designing a strain, as they can limit the interpretation of how specific tau modifications contribute to neurodegeneration.

To this end, the field has recently begun more carefully exploring how tau expression levels impact phenotypic outputs, with several papers, including recent papers from our lab, addressing this question in some depth ^10, 22, 45, 46^. The advent of single-copy gene insertion techniques has facilitated this process in *C. elegans*, but it also comes with limitations. In one of our previous studies, we generated and assessed single-copy, pan-neuronal T231E animals and found that touch responsiveness was selectively impacted ^10^; however, many commonly reported phenotypes were not present in our model, raising the concern that single-copy expression under the UPN promoter, while equal in all neurons, may be less than physiologically relevant.

To circumvent these caveats, here we generated stable transgenic lines using MosTI injection ^16^, which facilitates integrating multiple copies of a transgene into a safe-harbor loci as well as prevents mosaic expression and off-target effects from genomic integration (**Fig. 1A**). Most importantly, this approach – which we have termed “tunable tau”, allowed us to create a repertoire of stable transgenics with a spectrum of tau expression levels, from near single copy to high, albeit not to levels commonly observed in conventional extrachromosomal array-based approaches. We hope that this unique model will meld the advantages of the single-copy and overexpression approaches, adding a level of nuance to our ability to deconvolve dose dependent phenotypes from those that are more sensitive to disease-associated tau mutants. Using these strains, which possess nearly equivalent levels of tau in each individual neuron and are well-matched between wild-type T4 and T231E pairs, allowed us to clearly demonstrate that T231E tau can selectively impact specific behaviors and elicit unique mitochondrial unfolded protein responses. The specificity of the phenotypes to T231E tau is further supported by the fact that depletion through AID rescues dysfunction. Thus, we conclude that our tunable tau model provides a powerful platform for investigating the toxic potential of early-disease associated tau PTMs, as the level of control it provides allows for nuanced interrogation of dose-dependence, age-dependence, PTM specificity, and persistence of elicited phenotypes.

Disentangling the role that tau plays in normal physiology has been tricky, and the role it plays in AD pathology has eluded researchers equally so. Genes which are tied to increased AD risk are commonly assessed for insight into the cellular machinery that is critical for disease progression. On a cellular level, mutations in presenilins (PSEN1 and PSEN2) predispose patients to familial AD development and recent research has demonstrated that mutated presenilin leads to the accumulation of dysfunctional lysosomes ^47^. Of note, the laboratory of Kenneth Norman has extensively interrogated the neuronal consequences of PSEN mutations using the *C. elegans* orthologue *sel-12*, which also accumulates dysfunctional neuronal lysosomes when *sel-12* function is impaired ^48^. Interestingly, these mutants were shown to retain increased levels of mitochondrial calcium in their neurons, and lysosomal defects seen in the *sel-12* mutant background were largely rescued by limiting calcium uptake via mutation of mitochondrial calcium uniporter *mcu-1*. In a separate study, they observed that *sel-12* mutants possessed increased surface area within their mitochondria to endoplasmic reticulum contacts, suggesting that presenilin regulates organelle organization and communication ^49^. These results in *C. elegans,* in conjunction with observations of mitochondrial impairment in human AD patients, highlight the intersection of mitochondrial and neuronal function as a promising axis for sub-cellular pathology in AD progression.

We have previously demonstrated that single copy expression of T231E in touch neurons selectively suppressed stress-induced mitophagy in *C. elegans* mechanosensory neurons ^30, 33^. While we did not measure mitophagy in the multicopy strains due to confounds of spectral overlap with the reporter and pan-neuronal expression of tau, these previous results motivated us to ask whether stimulating mitophagy might suppress the multicopy phenotypes. We chose to focus on celastrol, a natural product and bioactive chemical compound which is a potent stimulator of mitophagy and is in wide use in preclinical trials ^31^. We found that inducing mitophagy through celastrol completely rescued both touch and thrashing deficits observed in our E7 strain (**Fig. 4A and 4B),** consistent with others’ observations on the benefits of stimulating mitophagy in neurodegenerative disease models ^24, 50, 51^.

*Dct-1* is the *C. elegans* homolog of mammalian BNIP3 and BNIP3L/NIX and is required for stress-induced mitophagy in *C. elegans* ^52^. Initially, we thought that genetic ablation of *dct-1* might prevent the beneficial effects of celastrol. To our surprise, however, loss of *dct-1* suppressed E7 phenotypes independent of celastrol (**Fig. 4C and 4D**). We consider there to be two reasonable possibilities. The first is that E7 acts through *dct-1* to cause neuronal dysfunction. However, it’s unlikely that E7 simply prevents DCT-1 activity, or the *dct-1(lf)* allele would be expected to phenocopy (or even exceed) E7 – and its loss does not impact either touch or thrashing behaviors (**Fig. 4**). The second and more likely possibility is that the loss of *dct-1* triggers a compensatory mechanism that protects against E7.

Our interest in the impact of compensatory mechanisms was further stimulated by finding that E7 animals were more resistant to ROS stress and lived longer than N2 controls (**Fig. 5**). This suggested that perhaps tau was perceived as a stress and itself induced a form of compensatory response. We suspected that adaptive stress responses may be involved in tau’s toxicity, as we have previously shown that a subtle but detectable cell autonomous UPRmt was present in touch neurons expressing T231E ^11^. Here, we used qRT-PCR to examine expression levels for select UPRmt genes shown to be highly upregulated in *sod-2* mutants ^53^, an approach that was not feasible in strains where tau expression was limited to six touch neurons ^11^. These results suggested frank activation of the UPRmt, though we did not assess cell autonomy (**Fig. 5**), and track with the observation that a mild activation of the UPRmt can extend lifespan and confer stress resistance to *C. elegans* ^54^.

There has been some recent work in other organisms on the topic of how the UPRmt can become maladaptive when chronically activated and may contribute to age-dependent diseases. Shpilka et al. demonstrated that the UPRmt plays a critical role in development, as it links mitochondrial biogenesis to nutrient sensation through mTORC1 activity. In short, their work showed that high mTORC1 activity levels during development lead to protein accumulation within mitochondria, which excludes ATFS-1 and promotes mitochondrial expansion ^55^. Another study demonstrated that nutrient sensation through Fox0/DAF-16 signaling promotes the accumulation of mutated mtDNA, similar to chronic ATFS-1 activity ^56^. These observations that UPRmt activity is tightly linked with mitochondrial homeostasis are not limited to *C. elegans*, as the accumulation of mtDNA defects within mice was shown to activate mTORC1 and subsequently trigger the UPRmt in mice ^57^. In human hematopoietic stem cells, a regulatory branch of the UPRmt mediated by SIRT7 and NRF1 and coupled to energy sensation was shown to modulate stem cell quiescence ^58, 59^. These studies identify one potential mechanism through which UPRmt signaling can become maladaptive: sustained stress can induce a heteroplasmic drift of mtDNA, providing the foundation for chronic UPRmt activation through cyclical nutrient sensing, mTORC1, and UPRmt positive feedback. Combined with neurodegenerative mutants that selectively inhibit mitophagy, this provides the foundations for a regulatory loop through which simple mitochondrial perturbations can coalesce into permanent shifts in mitochondrial homeostasis. Though still not completely understood, these shifts may create a cellular context that is tolerable at first but becomes neurodegenerative with age, when decreased mitochondrial function and impacted proteostasis are common. It has previously been reported that proteins associated with neurodegenerative diseases can become more neurotoxic when the UPRmt is chronically active ^60^. Given that disrupted UPRmt and mitochondrial homeostasis are commonly reported in AD neurons ^61^, our finding that the loss of *atfs-1* suppresses neuronal dysfunction in E7 animals demonstrates that the interplay between toxic tau epitopes and UPRmt signaling causes neuronal dysfunction. While we did not observe any neuronal loss in in our model, the fundamental mitochondrial machinery impacted by this interaction may become neurodegenerative with age or further insult and warrants additional research.

Interestingly, we found differential expression between T7 and E7 animals of several targets, indicating that the specific stress responses may be similar in terms of canonical UPRmt signaling yet unique in arms of the UPRmt response that are not as well studied (**Fig. 5**). This is relevant because E7 selectively exhibited phenotypes not observed in T7 and loss of *atfs-1*, central to the UPRmt, could suppress these phenotypes (**Fig. 5**). Specifically, *hrg-9* is uniquely upregulated in E7 animals. While *hrg-9*, whose human orthologue is TANGO2, is implicated in metabolic crisis disorders, its exact function and relevance to mitochondrial stress management is still being investigated ^39, 62^. Some reports show that loss of function in *hrg-9* elicit mitochondrial stress by preventing haem export from lysosome related organelles and subsequent maturation at mitochondria, leading to deficiencies in energy production and electron transport in the TCA cycle – loss of TANGO2 is even lethal in zebrafish ^39^. A recent study further showed that *hrg-9* responds to oxidative stress and likely helps to regulate mitochondrial function during crisis ^62^. Specifically, *hrg-9* knock out in *C. elegans* leads to defects in survival, locomotion, feeding, and brood size ^62^. Data from human patients and models of TANGO2 disorder show that vitamin B supplementation can lower the risk of crisis, suggesting that CoA synthesis or scavenging may be a critical metabolic hub through which *hrg-9* acts to mitigate mitochondrial stress ^62^. While it is interesting to speculate that T231E’s toxicity may be regulated through *hrg-9*, the effect of *hrg-9* overexpression is unknown. Additionally, *hrg-9* is predominantly expressed in the *C. elegans* intestine ^39^, which indirectly suggests that pan-neuronal expression of T231E elicits cell non-autonomous signaling in the rest of the animal. It also is likely that *hrg-9* is not the only UPRmt gene that is differentially regulated by T7 and E7, but the future identification of these differentially expressed genes may contribute to our understanding of why phenotypic differences occur between these strains. Ultimately, the goal would be to understand the relative impact of mitochondrial dysfunction versus dysregulation of adaptive signaling pathways on disease severity and progression.

In summary, we generated *C. elegans* models which allow us to tune pan-neuronal tau expression and assess phenotypic consequences as a function of protein expression level as well as their selectivity for a phosphomimetic tau variant (T231E) with an unprecedented level of control. Our results in aggregate, along with previous studies from our lab and others, demonstrate that early-disease tau phosphorylation events like pT231 are causative for neuronal disfunction and perturb mitochondrial quality control (MQC). Having a spectrum of strains with different tau expression levels facilitates studies of selective vulnerability, both at the level of neuron subpopulations’ susceptibility to tau induced dysfunction and molecular pathways underlying the effect of disease relevant modifications to tau. These observations support the general notion that MQC could be a critical event in early-AD development, and that therapies which target early-disease associated tau variants hold therapeutic promise.

## Acknowledgements

The authors would like to acknowledge BioRender and GraphPad Prism for aid in figure preparation.

## Author Contributions

Carroll, T. - Conceptualization, Investigation, Data Curation, Formal Analysis, Writing – Original Draft, Writing – Review and Editing, Visualization. Pfendler, D. – Conceptualization, Investigation, Data Curation. Alhaj Arhayem, H. – Investigation, Data Curation. Thoma, R. – Investigation. Müller-Eigner, A. – Investigation. Straut, A. – Investigation. Nehrke, K. and Johnson, G.V.W. – Conceptualization, Writing – Review and Editing, Funding Acquisition, Project Administration, Supervision.

## Author Approval

All authors have reviewed and approved the manuscript as is, and this article has not been accepted or published elsewhere.

## Ethical Considerations

Not applicable.

## Consent to Participate

Not applicable.

## Consent for Publication

Not applicable.

## Data Availability Statement

The data supporting the findings of this study are available within the article and/or its supplemental material.

---

**Supplemental Figure 1.**
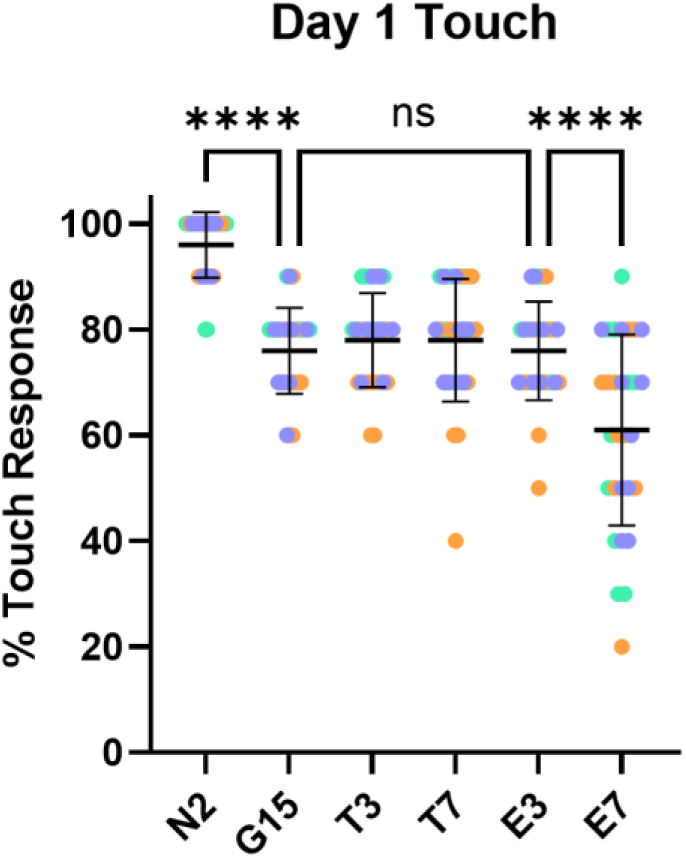
Touch Responsiveness of Tunable Tau Strains. The plots show mean ± SD of touch responsiveness in each strain at day 1 of adulthood, with each point representing an individual animal. N = 30 animals from three biological replicates. Each biological replicate is denoted by a unique color. Statistical analysis was performed using a one-way ANOVA followed by Tukey-corrected multiple comparisons. F = 28.47. DF = 209.

**Supplemental Figure 2.**
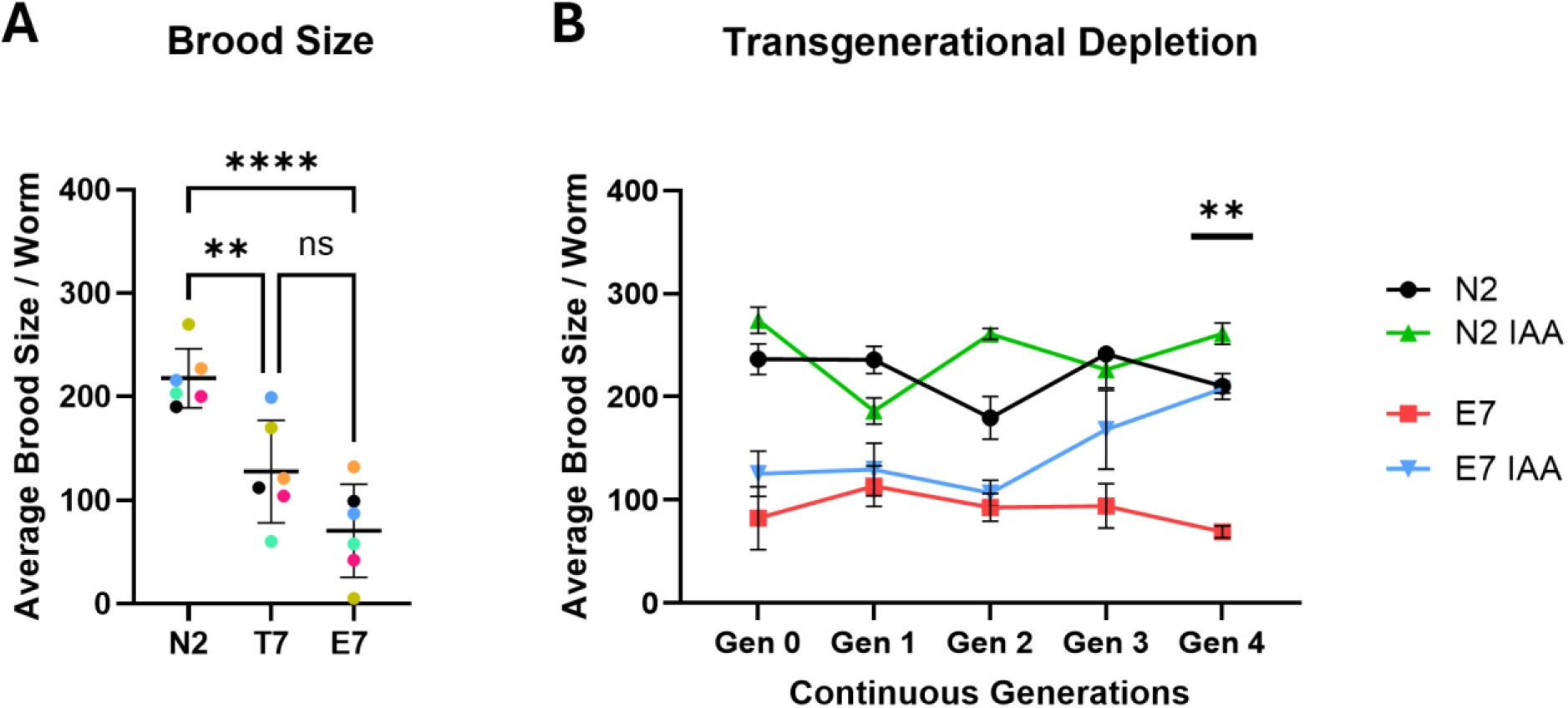
High-copy Tau Expression Causes Transgenerational Brood-size Deficits. A) Quantification of the total number of embryos laid from 3 animals over their entire lifespan. N = 6 biological replicates. Statistical analysis was performed using a one-way ANOVA followed by Tukey-corrected multiple comparisons. F = 18.68. DF = 17. B) Tracking of average brood size per worm for N2 and E7 animals over multiple generations on untreated or 4mM IAA plates to continuously deplete tau. N = 3-6 biological replicates. Reported values are mean ± SEM and reported significance is from the comparison between E7 and E7 IAA samples at Generation 4 using a three variable mixed-effects analysis followed by Tukey-corrected multiple comparisons. P < 0.0001 for IAA treatment as a variable. F = 29.40. DF = 37.

**Supplementary Table 1.**
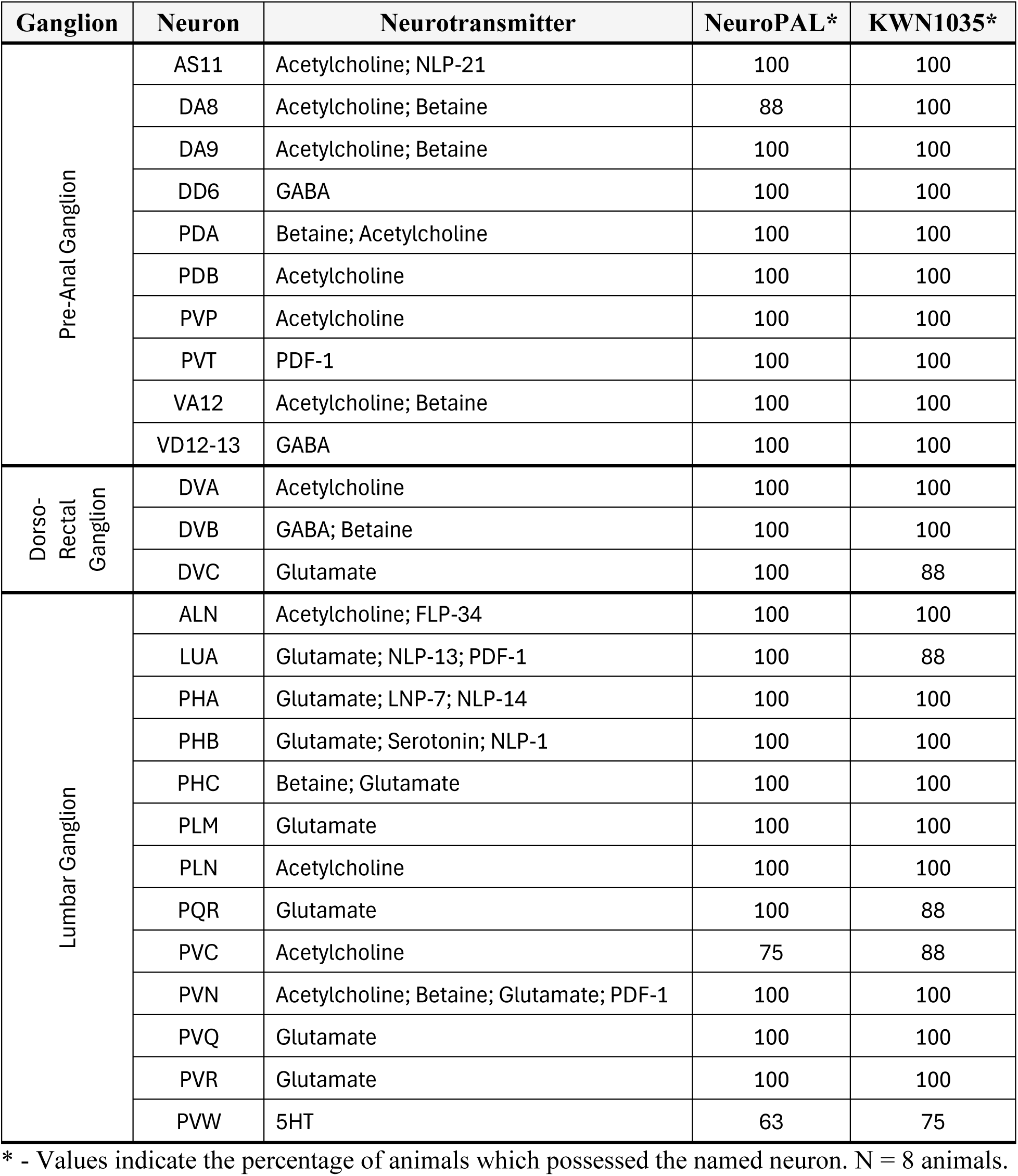

